# Colon Innervating TRPA1 Positive Nociceptors Influence Mucosal Health In Mice

**DOI:** 10.1101/2021.10.17.464752

**Authors:** Vibhu Kumar, Vijay Kumar, Kirti Devi, Ajay Kumar, Rehan Khan, Ravindra Pal Singh, Sivasubramanian Rajarammohan, Kanthi Kiran Kondepudi, Kanwaljit Chopra, Mahendra Bishnoi

**Author notes:** **Corresponding authors: Mahendra Bishnoi, PhD**, Scientist E, TR(i)P for Health Laboratory, Department of Food and Nutritional Biotechnology, National Agri-Food Biotechnology Institute (NABI), Knowledge City-Sector 81, SAS Nagar, Punjab 140306, India., **Co-corresponding authors Kanwaljit Chopra, PhD**, Professor, Pharmacology Division, University Institute of Pharmaceutical Sciences, Panjab University, Sector-14, Chandigarh, 160014, India; **Kanthi Kiran Kondepudi, PhD**, Scientist E, Department of Food and Nutritional Biotechnology, National Agri-Food Biotechnology Institute (NABI), Knowledge City-Sector 81, SAS Nagar, Punjab 140306, India.

## Abstract

**Introduction:** Transient receptor potential ankyrin-1 positive (TRPA1+ve) nociceptors, primarily present as peptidergic neuronal afferents in the colon are sensors of disturbance in lower gastrointestinal tract including pain induced by different pathologies. Their therapeutic role in the alleviation of chronic pain (receptor antagonism and receptor desensitization) associated with inflammatory bowel diseases (IBD) is reported. However, there is limited literature available about their role in formation and sustenance of the mucosal layer, and its interaction with host physiology as well as luminal microbial community. The aim of this study focuses on the effects of nociceptive TRPA1 channel desensitization on colonic mucus production and gut health.

**Methods:** TRPA1+ve nociceptors were desensitized by rectal administration of capsazepine. Ileum, colon was harvested and cecum content was collected. We performed morphological/histological analysis, gut permeability alteration, gene expression changes, colon metabolite profiling, and gut microbial abundance in these animals.

**Results:** We found that presence of TRPA1-positive nociceptors is required for mucus layer integrity, using an intra-rectal capsazepine-induced TRPA1 desensitization model. Desensitization of TRPA1 positive nociceptors resulted in damaged mucosal lining, resultant increase in gut permeability and altered transcriptional profile of genes for goblet cell markers, mucus regulation, immune response and tight junction proteins. The damage to mucosal lining prevented its role in enterosyne (short chain fatty acids) actions.

**Conclusion:** These results suggest that caution may need to be exercised before employing TRPA1 desensitization as a therapeutic option to alleviate pain caused due to IBD.

## 1. Introduction

Sensory neurons innervating the gastrointestinal tract play a significant role in IBD^1^. Release of neuropeptides, calcitonin gene-related peptide (CGRP) and substance P (SP), from sensory neurons induce vasodilation and activation of immune cells^2^. Transient receptor potential (TRP) ion channels, particularly Vanilloid-1 (V1) and Ankyrin-1 (A1), have significant expression in sensory neurons and are reported to be involved for symptoms of IBD^1, 3^ or colonic distension pain in rodents^4^ In the rodent model of dextran sodium sulfate (DSS)-induced colitis, neuronal TRPA1 sensitization in the colon resulted in SP release that contributes to inflammation^1^. In same study, TRPA1 agonist 2,4,6-trinitrobenzene-sulfonic-acid (TNBS), was used for induction of colitis in mice^1^. Evidence exists on the use of TRPA1 agonists, AITC and formalin, to induce colitis in rodents^5, 6^. Targeted TRPA1 channel inhibition can be a therapeutic approach for alleviation of IBD associated pain^1, 7, 8^.

The intestinal mucus layer protects against mechanical, chemical and biological attacks and contributes to the maintenance of intestinal homeostasis^9^ as an interplay between host and microbe. Alterations like decrease in mucin layer can induce gut dysfunction^10^. There is limited literature directly associating TRPA1 channel with maintenance of mucus secretion in the gut. A recent study revealed that TRPA1 agonist cinnamaldehyde induced HCO^3-^ secretion in colon, which is essential for mucus release^11^. Recently published work from our laboratory demonstrated that systemic TRPV1 ablation leads to impaired mucus secretion and causes dysbiosis in the gut^12^. Given the importance of TRPA1 and TRPV1 in gut mucin-immune axis, we hypothesize that the strategies of blocking (with antagonists) or desensitizing sensory nociceptive neuronal TRPV1 and TRPA1 for the treatment of IBD might impair colonic mucin homeostasis, leading to increased gut permeability and dysbiosis.

Recently efforts to classify colonic sensory neurons using single cell RNA sequencing analysis suggest a high percentage of TRPA1^+ve^ sensory peptidergic neurons^13^. Sensing the significance of TRPA1 (high expression in colonic sensory neurons and co-expression with TRPV1), the present study was designed to understand the role of TRPA1^+ve^ nociceptive sensory neurons in gut mucosal health using rectal capsazepine-induced systemic TRPA1 desensitization model in mice. We further sought to explore whether these nociceptive neurons have a modulatory role in the action of SCFAs, a common nutrient and microbe derived enterosyne in the colon. These findings will serve to inform considerations on unwanted effects of TRPA1 (or TRPV1) desensitization or blockade as a treatment strategy for IBD induced pain.

## 2. Results

### 2.1. Rectal administration of CPZ induced systemic desensitization of neuronal peptidergic nociceptive TRPA1 channels in DRGs

Rectal administration of CPZ significantly altered TRPA1 associated nociception as evident from reduced AITC-induced eye wipes (Fig. S1A) and elevated tail withdrawal latency time at 4°C (Fig. S1B) as compared to control animals. These effects were constant up to the end of the study (day-14). There was significant decrease in capsaicin induced eye wipes and significant increase in tail withdrawal latency at 42°C at day-14 as compared to control group (Fig. S1C). The representative figure from the study of Hockley *et al*., 2019^13^ shows that 49% of TRPA1 expressing colonic sensory neurons belong to mPeptidergic-type-b (mPEPb) class (Fig. S1D). CPZ treatment significantly reduced the expression levels of *Spp1, Trpa1, Th and Calca* genes (Fig. S1E) as compared to control animals, which specifically express in mPEPb class of colonic sensory neurons.

### 2.2. CPZ-induced desensitization of neuronal peptidergic TRPA1 negatively regulates mucus homeostasis

Histological analysis of colon at day 14 revealed that CPZ administration markedly reduced the thickness of the mucus layer in lumen of proximal and distal colon (Fig. 2A, B, C) as compared to control animals. In addition to this, in distal colon, a significant decrease in goblet cell numbers as well as alcian blue (AB) and high iron diamine (HID) stain intensity (as a measure of diminished sialomucins and sulfomucins) was observed after CPZ treatment (Fig. 2A). Ileum did not show any morphological changes. SEM analysis showed a major decrease in number of mucus extrusions (M) from goblet cells (GC) and crypts (C) of colon indicating reduced mucus release after CPZ treatment. Also, the mucosal architecture with smooth velvet appearance in control group was lost upon CPZ treatment (Fig. 2D, E). Substantiating these findings, expression levels of genes involved in goblet cell functioning-*Cdx2, Dll1, Foxa2, Tff3, Vamp8* and *Prkd1* were downregulated in distal colon following CPZ administration (Fig. 2F). Difference was observable in genes-*Dll4* and *Foxa3* but were statistically insignificant (Fig-2F). Genes involved in colonic mucus production–*Muc2* and *Muc3*, were repressed in distal colon of CPZ treated animals (Fig-2G). Alternatively, expression of *Muc15* was significantly elevated (Fig-2G). Another set of genes involved in immune response – *Nlrp3, Nod2* and *Tnfα* were significantly upregulated, while *Fcgbp* expression level was reduced with CPZ treatment (Fig-2H).

**Figure-1:**
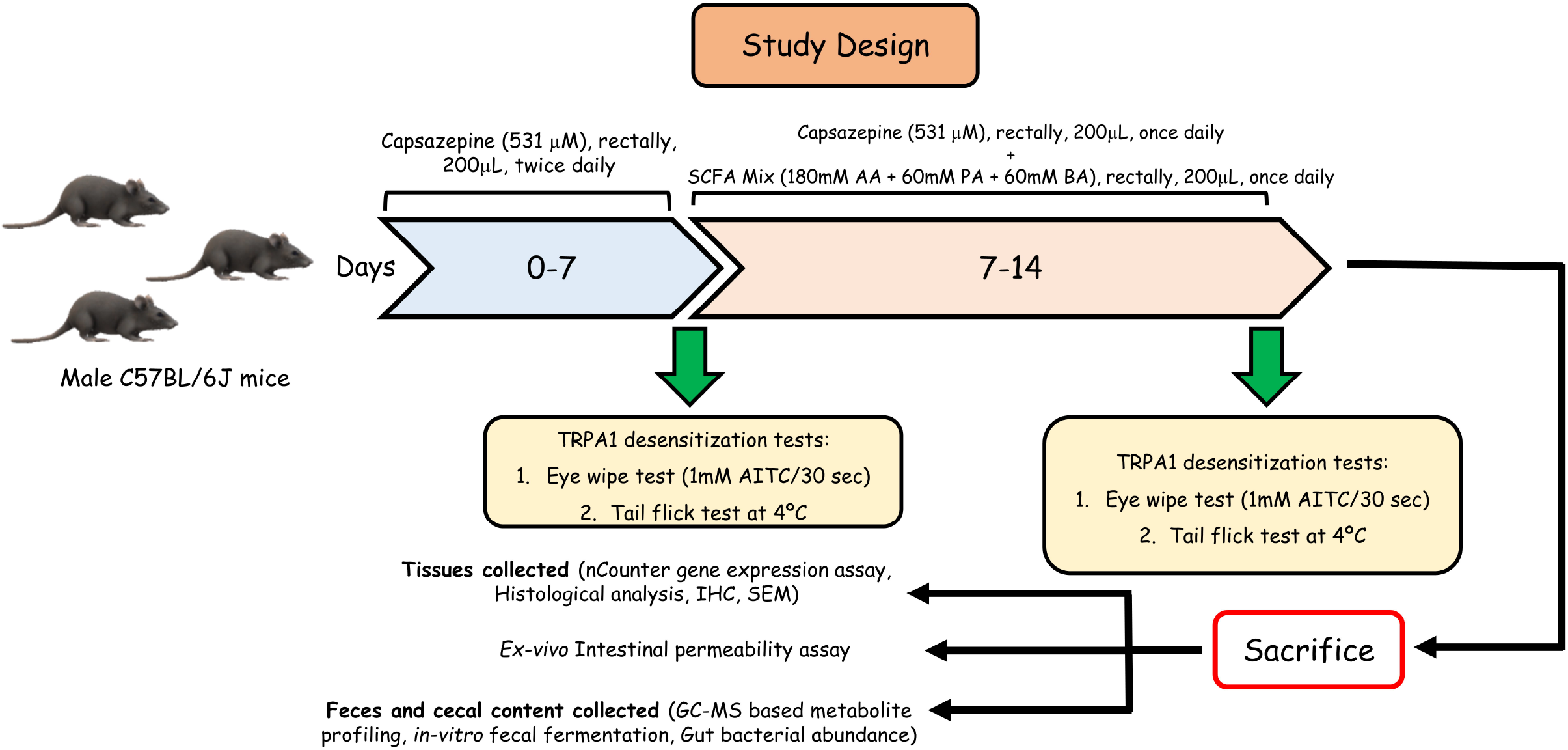
Animal experiment plan.

**Figure-2:**
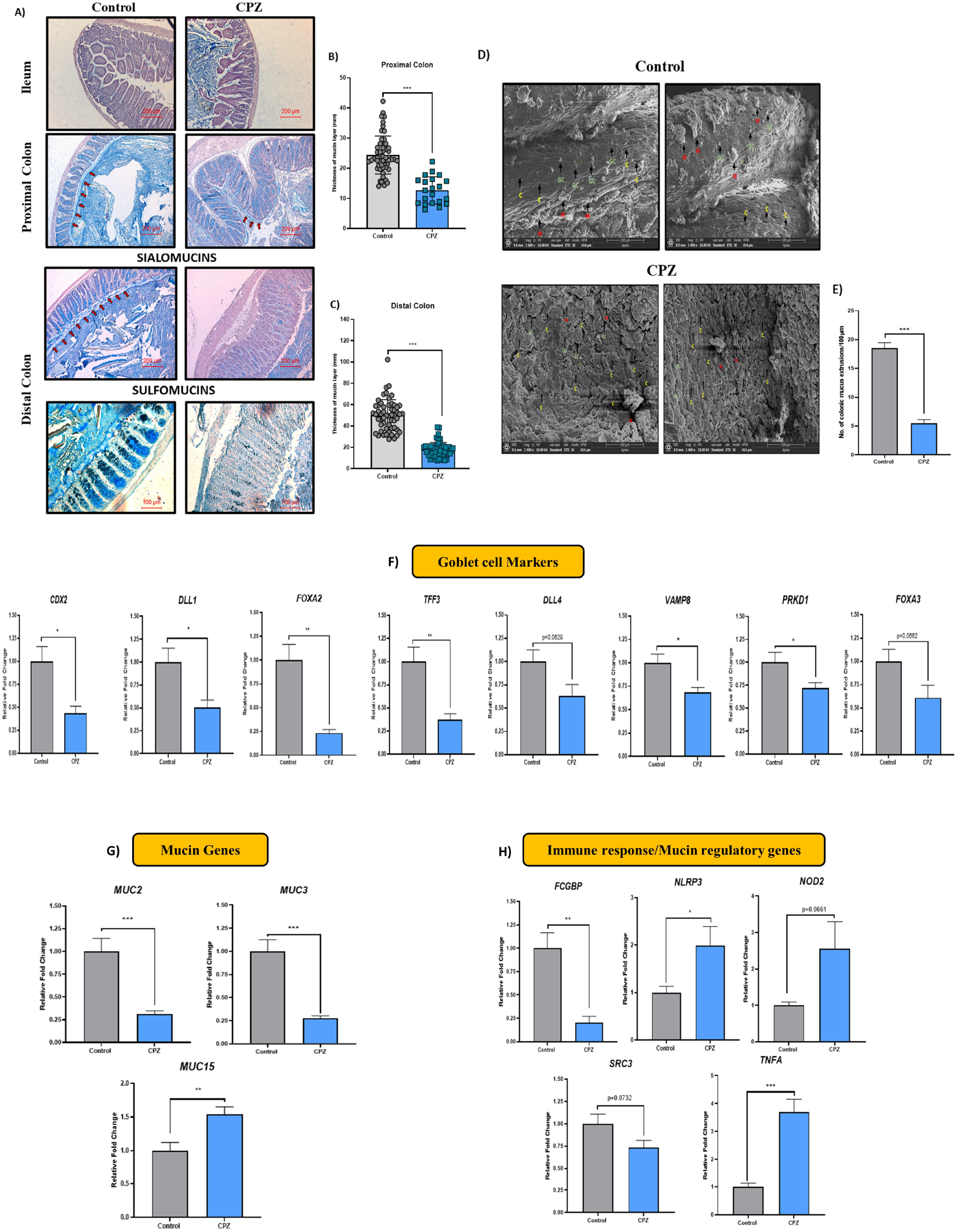
Effect of CPZ rectal administration on colonic mucus production. Histology representative images of A) Hematoxylin & Eosin – Alcian blue staining in ileum, proximal and distal colon (n=3), High iron diamine – Alcian blue staining in distal colon (n=3), B) Thickness of mucus layer in proximal colon, C) Thickness of mucus layer in distal colon. D) Representative images for SEM of colon tissues and E) Number of colonic mucus extrusion. Gene expression analysis for F) Goblet cell markers, G) Mucin genes and H) Immune response/mucus regulation in colon (n=6). Mice were divided into two groups - Control (administered vehicle rectally) and CPZ (administered 531μM CPZ rectally). Treatment was given for 2 weeks. After sacrifice, colon tissues were fixed in Carnoy’s solution, subjected to serial dehydration with gradient ethanol (25%, 50%, 75%, 90% and twice in 100%) 2 h each followed by clearing in xylene (1 h twice). The colon tissues were embedded in paraffin and 5μm sections were cut. For staining, the sections were deparaffinized in xylene (5 min, twice) followed by rehydration in gradient ethanol (100% twice, 90%, 70%, 50%, 25% and PBS, 2min each). The samples were treated with respective stains (H&E: 2min each, AB: 15min, HID: 24h) followed by washing (5min, thrice after each stain) in water. Sections were dehydrated, treated with xylene and mounted with DPX mounting medium. Microscopy was performed for imaging and mucus thickness was measured using Imagescope. For SEM, colon sections were cut open (lumen was exposed) and fixed in 2.5% glutaraldehyde. Following, the tissues were washed with PBS (15 min, thrice at 4°C). The tissues were post-fixed in 1%osmium tetroxide for 2 h and again washed with PBS (15 min, thrice at 4°C). The tissues were then serial dehydrated in acetone (30%, 50%, 70%, 90%, 95%, 100%, 30 min at 4 °C). The samples were air dried and spur-coated with gold (10 nm) and images were taken on scanning electron microscope. Colonic mucus extrusions (M) from goblet cells (GC) and crypts (C) of colon were counted using Imagescope. For gene expression studies, RNA was extracted from colon tissues and gene expression was performed for goblet cell markers, mucin genes and genes involved in immune response and mucus regulation using Nanostring nCounter multiplex gene expression assay. All data is represented as mean ± SEM. Intergroup variations were assessed using Student unpaired t-test. *p<0.05, **p<0.01, *** p<0.001 versus Control.

### 2.3. CPZ-induced TRPA1 desensitization increases intestinal permeability and compromises gut barrier function

Ex-vivo intestinal permeability was evaluated by time dependent fluorescent imaging. In this study, the diffused FITC loaded nanoparticle samples withdrawn at different time points were imaged and analyzed for membrane barrier function. We have observed intense fluorescent intensity in the CPZ group as compared with control group (Fig. 3A, 3B). Furthermore, the area under curve (AUC) in CPZ treated animals indicating a higher diffusion rate of FITC-loaded nanoparticles from the colon and suggested a compromised intestinal permeability (Fig. 3C). It has also been observed that the expression of Tight Junction Proteins (TJPs; Zona occludens-1 (*ZO-1*), Occludin (*OCC*) and Claudin-1 (*CLDN-1*)) in colon sections was downregulated in CPZ group as compared to the control group as indicated by immunohistochemical evaluation (Fig. 3D, E, F). Gene expression studies also indicated a similar trend of reduction in expression of *Cdh, Cldn-1, Zo-1* and *Occ* in colon of CPZ treated animals as compared to control group (Fig. 3G).

**Figure-3:**
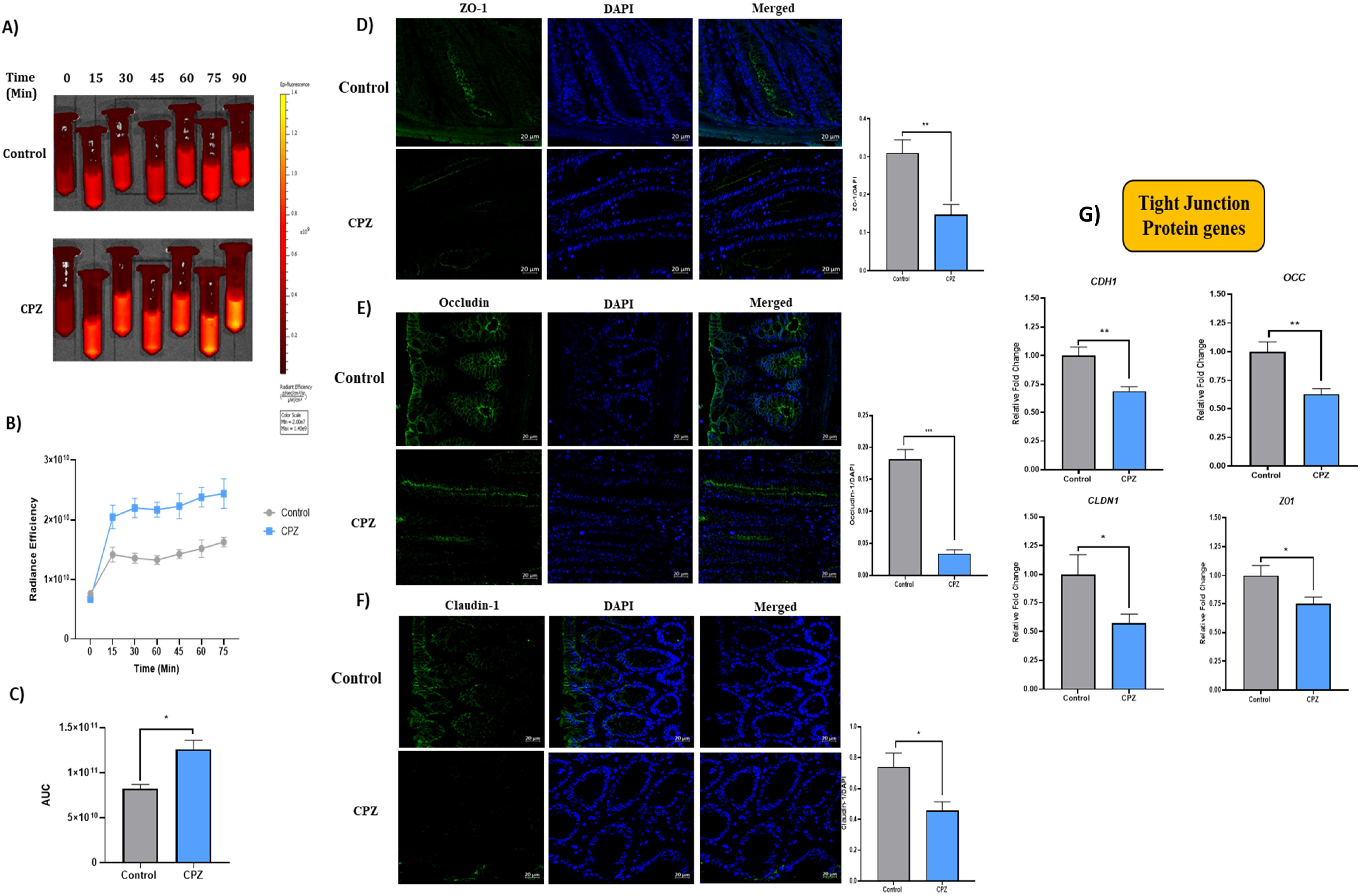
Effect of CPZ rectal administration on intestinal permeability and tight junction proteins. A) *Exvivo* intestinal permeability – representative images for release of FITC-loaded nanoparticles from colon at different time points (n=3), B) Time-based plot for radiance efficiency and C) AUC of the radiance efficiency. Representative images for immunofluorescence in colon tissues for expression of - D) Zona occludens-1 (*ZO-1*), E) Occludin (*OCC*) and F) Claudin-1 (*CLDN-1*). Fluorescence intensity were measured using ImageJ and data is presented after normalization with DAPI. G) Gene expression analysis for tight junction proteins. Mice were divided into two groups – Control (administered vehicle rectally) and CPZ (administered 531μM CPZ rectally). Treatment was given for 2 weeks. After sacrifice, colon tissues (n=3) were fixed in carnoy fixative, paraffin embedded moulds were prepared and 5μm sections were obtained. The sections were subjected to IHC with antigen retrieval using citrate buffer pH 6.0 (90°C, 60 min) followed by blocking in 2.5% goat serum in PBS-Tween20, overnight incubation in primary antibodies, secondary antibody treatment for 2 h and DAPI for 30-60s. High resolution images were obtained using confocal microscope and florescence intensity were measured using ImageJ. For gene expression studies, RNA was extracted from colon tissues and gene expression was performed for tight junction proteins using Nanostring nCounter multiplex gene expression assay. All data is represented as mean ± SEM. Intergroup variations were assessed using Student unpaired t-test. *p<0.05, **p<0.01, *** p<0.001 versus Control.

### 2.4. Desensitization of nociceptive peptidergic TRPA1 alters mucus associated cecal metabolites and mucin glycosylation enzymes

GC-MS based metabolome of cecal content displayed significantly lower levels of mucus related glycans (fructose, ribose, xylose, xylulose, glucose and arabinose) and proline, one of the core amino acids of PTS domain in mucin for its structural integrity (Fig. S2A). In addition, gene expression for enzymes involved in mucus glycosylation – *B4galnt-2, C3gnt-1, St3gal4, Galnt-1, Galnt-3* were also significantly downregulated in CPZ group as compared to control (Fig. S2B).

### 2.5. Rectal administration of CPZ induced gut microbiota changes

CPZ treatment increased the abundance of Gram^+ve^ bacteria - *Roseburia, Lactobacillus, Akkermansia, Fecalibacterium, Eubacteria, Ruminococci, Butyricicoccus pullicaecorum, Clostridium coccoides, Anaerostipes butyraticus*, Lachnospiraceae and Firmicutes significantly (Fig. 4A, B). However, abundance of Gram^-ve^ bacteria–*Butyrivibrio fibrisolvens, Bacteroides fragilis, Prevotella copri*, Bacteroidetes, Bacteroides and Fusobacteria remained unaltered with CPZ treatment. (Fig. 4C). Only Cronobacteria was increased in significant manner as compared to control group (Fig. 4C). Firmicutes-Bacteroides ratio was significantly higher in CPZ treated animals (Fig. 4D). Further, to evaluate host-independent effects of CPZ on gut microbiota, fresh feces from normal mice were subjected to *in-vitro* fermentation in presence of CPZ. qRT-PCR analysis revealed that abundance of Gram^-ve^ bacteria increased – *Butyrivibrio fibrisolvens, Prevotella, Bacteroidetes, Fusobacteria, Bacteroides and Cronobacteria* (Fig-S3A). Out of Gram^+ve^ bacteria, *Firmicutes, Roseburia and Lactobacillus* were elevated with CPZ treatment as compared to control (Fig-S3B). No effect was observed on other Gram^+ve^ genera–*Akkermansia, Fecalibacterium, Eubacteria, Ruminococci and Anaerostipes butyraticus* in presence of CPZ.

**Figure-4:**
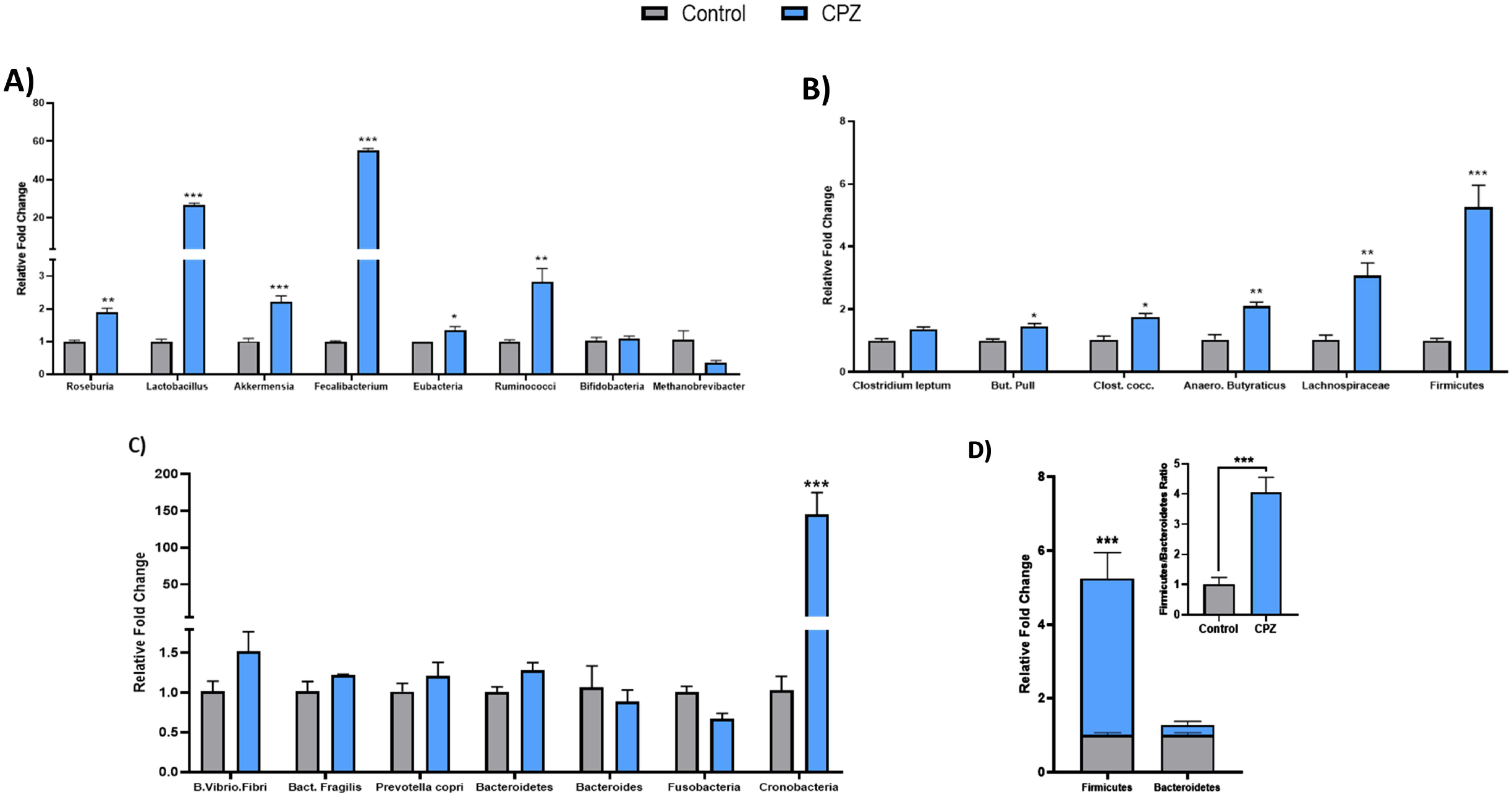
Effect of CPZ rectal administration on gut microbiota. A), B) Relative bacterial abundances of Gram +ve bacteria in cecum, C) Relative bacterial abundances of Gram -ve bacteria in cecum and D) Relative abundance of firmicutes and bacteroides in cecum. Mice were divided into two groups – Control (administered vehicle rectally) and CPZ (administered 531μM CPZ rectally). Treatment was given for 2 weeks. After sacrifice, the cecum content was collected, weighed and bacterial DNA was isolated using commercial kit. The DNA quantity was measured and quality was assessed with agarose gel electrophoresis. The DNA was subjected to qRT-PCR based changes in gene expression of various gut bacteria. The data was normalized with total bacteria as internal control and presented in the form of relative fold change. All data is represented as mean ± SEM. Intergroup variations were assessed using Student unpaired t-test. *p<0.05, **p<0.01, *** p<0.001 versus Control.

### 2.6. Rectal co-administration of SCFA mix fails to prevent CPZ - induced compromise in colonic mucin homeostasis

Rectal administration of SCFA mix *per se* increased the thickness of the mucus layer, as well as sialomucins and sulfomucins content in goblet cells. However, its co-administration with CPZ is unable to prevent the CPZ-mediated reduction in goblet cells differentiation and mucus production (Fig. 5A, B, C). Colon tissues from SCFA *per se* treated mice displayed a significant increase in mucus extrusions (M) from goblet cells (GC) and crypts (C) of colon, but minimal effect was observed upon their co-administration with CPZ. Also, SCFA co-administration was unable to reverse the CPZ-induced morphological alteration in mucosal architecture (Fig. 5D, E). Further, the SCFA mix intervention increased the expression of TJPs – *ZO-1, OCC* and *CLDN-1*, but CPZ-induced downregulation in these proteins was unaffected with SCFA mix co-administration (Fig. 5F, G, H). Similarly, qRT-PCR results of *Cldn-1* and *Zo-1* displayed similar trend but were statistically non-significant (Fig. 5I).

**Figure-5:**
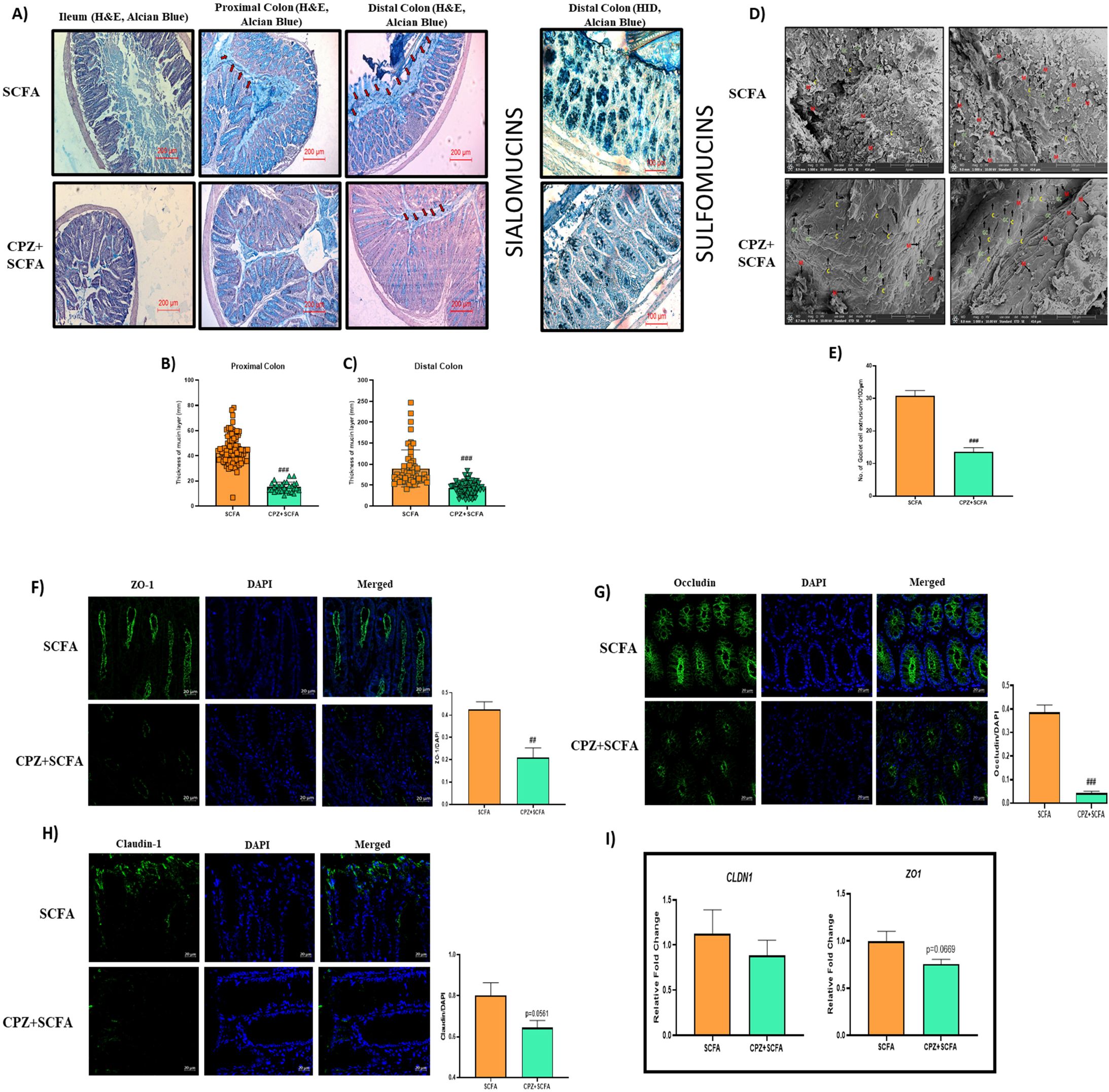
Effect of rectal co-administration of SCFAs on CPZ-induced alterations in colonic mucus production and tight junction proteins. Histology representative images of A) Hematoxylin & Eosin – Alcian blue staining in ileum, proximal and distal colon (n=3), High iron diamine – Alcian blue staining in distal colon (n=3), B) Thickness of mucus layer in proximal colon, C) Thickness of mucus layer in distal colon. D) Representative images for SEM of colon tissues and E) Number of colonic mucus extrusion. Representative images for immunofluorescence in colon tissues for expression of - F) Zona occludens-1 (*ZO-1*), G) Occludin (*OCC*) and H) Claudin-1 (*CLDN-1*) along with their fluorescence intensities. Fluorescence intensity were measured using ImageJ and data is presented after normalization with DAPI. I) Gene expression analysis for tight junction proteins. For Histology, Mice were divided into two groups (n=6): SCFA (treated with rectal administration 200μL SCFA mix [180mM Acetic acid + 60mM Propionic acid + 60mM Butyric acid for 14 days]) and CPZ+SCFA (initially treated with 531μM CPZ twice daily for 7 days followed by once daily dose of 531μM CPZ along with rectal administration of SCFA mix for next 7 days). For staining, the sections were deparaffinized in xylene (5 min, twice) followed by rehydration in gradient ethanol (100% twice, 90%, 70%, 50%, 25% and PBS, 2min each). The samples were treated with respective stains (H&E: 2min each, AB: 15min, HID: 24h) followed by washing (5min, thrice after each stain) in water. Sections were dehydrated, treated with xylene and mounted with DPX mounting medium. Microscopy was performed for imaging and mucus thickness was measured using Imagescope. For SEM, colon sections were cut open (lumen was exposed) and fixed in 2.5% glutaraldehyde. Following, the tissues were washed with PBS (15 min, thrice at 4°C). The tissues were postfixed in 1%osmium tetroxide for 2 h and again washed with PBS (15 min, thrice at 4°C). The tissues were then serial dehydrated in acetone (30%, 50%, 70%, 90%, 95%, 100%, 30 min at 4 °C). The samples were air dried and spur-coated with gold (10 nm) and images were taken on scanning electron microscope. Colonic mucus extrusions (M) from goblet cells (GC) and crypts (C) of colon were counted using Imagescope. For IHC, colon tissues (n=3) were fixed in carnoy fixative, paraffin embedded moulds were prepared and 5μm sections were obtained. The sections were subjected to IHC with antigen retrieval using citrate buffer pH 6.0 (90°C, 60 min) followed by blocking in 2.5% goat serum in PBS-Tween20, overnight incubation in primary antibodies, secondary antibody treatment for 2 h and DAPI for 30-60s. High resolution images were obtained using confocal microscope and florescence intensity were measured using ImageJ. For gene expression, studies, RNA was extracted from colon tissues and gene expression was performed for tight junction proteins using Nanostring nCounter multiplex gene expression assay. All data is represented as mean ± SEM. Intergroup variations were assessed using Student unpaired t-test. *p<0.05, **p<0.01, *** p<0.001 versus SCFA.

**Figure-6:**
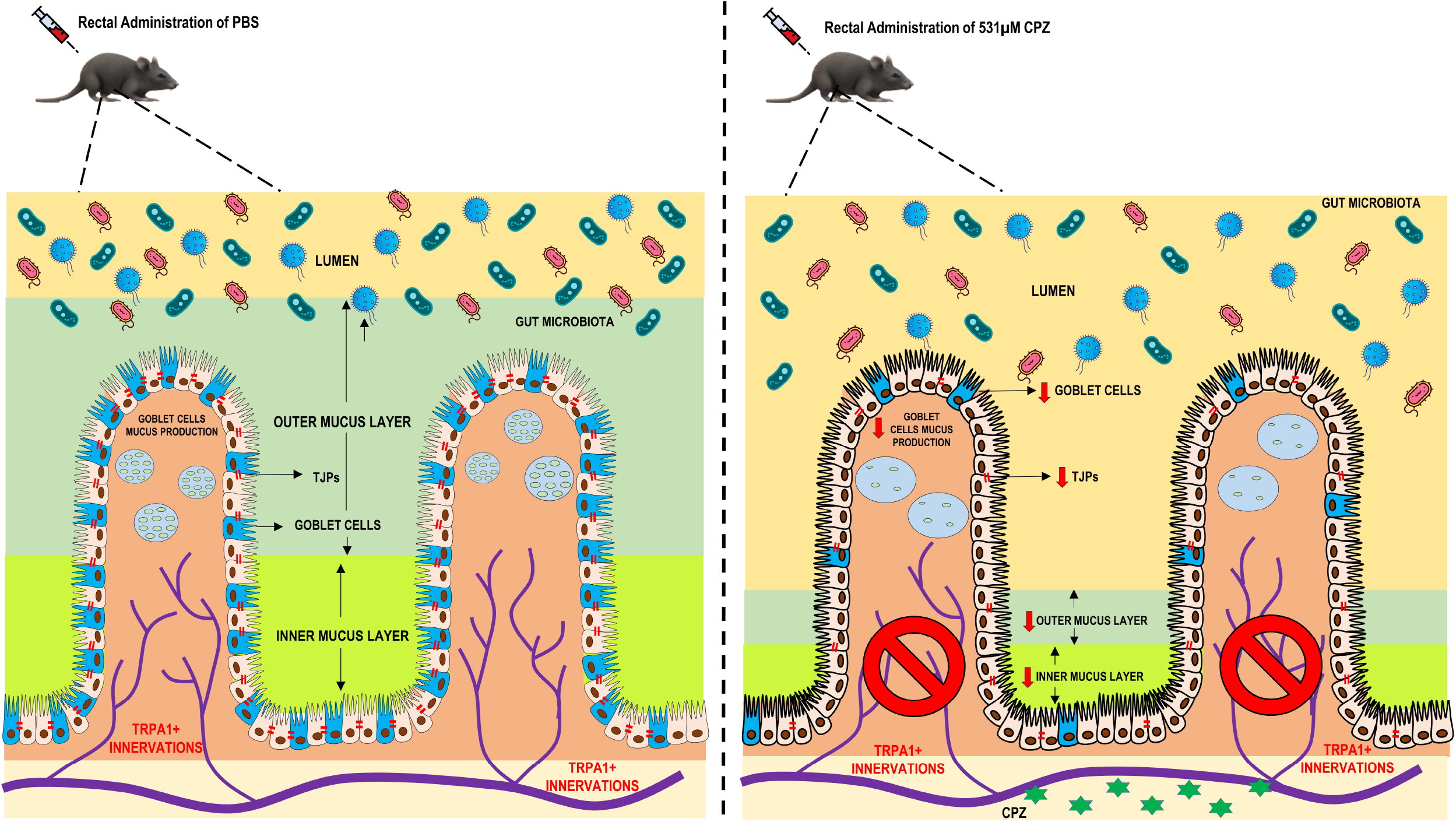
Representative image for overall effects of CPZ-induced desensitization of peptidergic nociceptive TRPA1 neurons on colonic mucus production and gut health.

### 2.7. Principal component and correlation analysis

Overall, the principal component (PC) analysis clearly showed that the CPZ group was distinct from the control group with respect to the gut microbiota, metabolites, and host gene expression (Figure-S4A, S4B and S4C). Additionally, most of parameters assessed displayed a negative correlation among themselves and other parameters between the Control and CPZ group (Figure-S4D). A similar pattern was observed in the PC and correlation analysis of the same parameters in SCFA and CPZ-SCFA group. The PC analysis for gut microbiota and metabolites showed higher dissimilarity between SCFA and CPZ-SCFA group (Figure S5A and S5B). Gene expression profiling however, showed only a slight dissimilarity between both of the groups (Figure S5C) indicating that CPZ mediated TRPA1 desensitization inhibits the beneficial effects of SCFAs in colon. In addition, the pattern in correlation matrix displayed a negative correlation in most of the parameters between both the groups (Figure S5D).

## 3. Discussion

Nociceptors, in particular peptidergic TRPV1 and TRPA1, regulate epithelial cell function (including goblet cells) owing to their involvement in neurogenic inflammation^8, 14^ There are numerous reports delineating their role in various gut-related diseases^1, 15, 16^. Release of these mediators and changes in mucus production in gut epithelial layer, in response to nociceptor activation have an ability to affect microbial population in a cyclic manner. Moreover, there is literature describing the reciprocal relationship between gut microbial population and its role in mucin secretion, which influences not only mucus layer formation but also its regulation^17, 18^. In our study, desensitization of TRPA1^+ve^ nociceptors caused loss of mucin layer, prominently in the colon. Both forms of mucin (sialomucin and sulfomucin) were found to be significantly decreased. Hence, we may say that functional absence of TRPA1^+ve^ nociceptors contributed, at least in part, to the damage of the mucosal layer, making hosts more susceptible to localized infection through colonization of harmful bacteria, food borne antigens and chemicals. However, the specific interactions and mechanisms are still infancy and an interesting area for further research. TRPA1 nociceptor desensitized mice had significantly enhanced gut permeability as studied by *ex-vivo* nanoparticle diffusion approach through colon. In desensitized animals both protein and gene expression for tight-junction proteins (TJP)s were significantly decreased suggesting both translational and transcriptional decrease in tight junction morphology. These findings are of significance because desensitization of nociceptive neurons prevented the SCFA effect in these animals^19, 20^, hence suggesting that intact mucin layer is important for the effects of healthy microbial metabolites in the host gut.

*Muc3* and *Muc2* genes were significantly decreased in desensitized animals. In studies on *Muc2* knockout animals, increased intestinal permeability has been shown to triggers a cascade of events leading to secondary organ inflammation and translocation of bacterial metabolites and bacteria *per se* leading to sepsis^21, 22^ Different goblet cell markers (*Cdx2, Dll1, Foxa2, Tff3, Dll4, Vamp8, Prkd1, and Foxa3*) and *Fcgbp* were significantly decreased in desensitized animals. Decrease in *Muc 3* (a major transmembrane mucin present on the N-terminal of extracellular mucin domain tip of the enterocyte microvilli) points to compromised cell protection (due to its role in densely glycosylated glycocalyx) and altered signal transduction pathways associated to microbe: host interactions^23^. In contrast, another cell surface associated with *Muc15* was significantly higher in desensitized animals and has role in carcinogenesis and protein metabolism^24^. Overall, alteration in levels of these mucin secreting and transmembrane genes is suggestive of compromised mucin homeostasis over all, making desensitized animals prone to disorders. This is reflected in the increase in the expression of genes related to immune response activation (*Nlrp3, Nod2, Tnfα*)^25, 26^.

Mucin layer is a source of nutrition and provider of attachment sites for gut microbes^27^. In the absence of mucin layer (decrease in glycosylation marker genes) in TRPA1 desensitized mice, colonic bacteria are deprived of both nutrients and their attachment sites forcing them to rely more on diet-extracted nutrition. A major part of diet comprises of plant dietary fiber, *i.e*. high molecular weight carbohydrates (polysaccharides). Metabolite analysis in cecal samples showed a decrease in the monomeric metabolites of these polysaccharides in TRPA1 desensitized animals indicating greater utilization of these by local microbiota as a nutrient source. During bacterial profiling in these samples, the increase in the (a) levels of Firmicutes and (b) ratio of Firmicutes/Bacteroidetes suggests increased extraction of energy from the dietary sources^28^. There was a significant increase in good commensal bacteria (*Lactobacillus, Fecalibacterium* and *Akkermensia etc*) in CPZ treated animals. We suggest this is an adaptive response to decrease in mucin layer, as we did not see any significant effect of capsazepine in individual groups of bacteria during *in-vitro* fecal batch fermentation experiments. Overall, the PC and correlation analysis indicate that (a) TRPA1 desensitization caused considerable changes in overall microbiota, metabolite and gene expression profile (b) prevented beneficial effects of SCFAs.

In conclusion, it is still not sufficiently clear whether side effects presented here and detrimental (mucous layer loss) or beneficial (adaptive response of increase in gut microbial ecosystem). These studies are of potential clinical applications as TRPA1/TRPV1 antagonism and desensitization are marked as promising therapeutic approaches for the alleviation of pain associated with IBD. Following our evidence presented here, we suggest further exploration into the role of nociceptors in mucin health and host: microbe interactions for better understanding of side effect profiles of these approaches before taking them to clinical practice.

## 4. Materials and Methods

### 4.1. Animals

Six-eight-week-old male C57BL/6J mice (25-30g), procured from IMTech Center for Animal Resources and Experimentation (iCARE; 55/GO/Re/Rc/Bi/Bt/S/99/CPCSEA), Chandigarh, India, and housed (temperature, 24 ± 2 °C; humidity, 55-65 %, 12h light-dark cycle) in Animal Experimentation Facility (2039/GO/RBi/S/18/CPCSEA), National Agri-Food Biotechnology Institute (NABI), Mohali, India. Animals were provided with water and normal pellet diet (Hylasco Biotechnology, India) *ad libitum*. Experimental protocol was approved (Approval number NABI/2039/CPCSEA/IAEC/2019/16) by Institutional Animal Ethics Committee (IAEC) of NABI. Committee for the Purpose of Control and Supervision on Experiments on Animals (CPCSEA) guidelines on the use and care of experimental animals were followed throughout the experiment.

### 4.2. Neuronal nociceptive TRPA1 channel desensitization, confirmation tests and SCFA mix treatment

After 1 week of acclimatization, animals were divided into 4 groups – Control, Capsazepine (CPZ), SCFA (Short chain fatty acids) and CPZ+SCFA (n = 6 each). Systemic desensitization of nociceptive peptidergic TRPA1 neurons in colonic dorsal root ganglia (DRG) was achieved by previously described method with slight modification^8^. Rectal administration of 531μM Capsazepine (CPZ) (200μL) (Cat. No. C191-25MG, Sigma-Aldrich) enemas was given to 12 animals twice daily for seven days for induction of desensitization followed by once daily administration for next seven days for its maintenance under brief isoflurane anesthesia. Control animals received 200μL PBS rectally as vehicle. Desensitization of nociceptive peptidergic TRPA1 neurons was confirmed by loss of nociceptive physiological responses after 7 days using tail flick test and eye wipe test. For tail flick test, tip of the tail of mice was dip in cold water (4°C) and time taken to withdraw the tail was noted as pain response (cut-off time of 30 s). Experiments were repeated 3 times/animal, at intervals of 5 min. For eye-wipe test, 1mM AITC solution (Cat. No. 377430; Sigma-Aldrich) was instilled into the eye of the animal and number of wipes was recorded for 30s. After 7 days, the animals receiving CPZ were divided in two groups – one group (CPZ+SCFA) received 200μL SCFA mix [180mM Acetic acid + 60mM Propionic acid + 60mM Butyric acid (neutralized to pH 6.5); Cat. No. A6283, 402907, B103500; Sigma-Aldrich, USA] rectally, once a day along with once daily dose of 200μL of 531μM CPZ rectal administration. The other group (CPZ) only received once daily administration of 200μL 531μM CPZ as maintenance dose for desensitization for next 7 days. Simultaneously, another set of animals were also receiving rectal administration of 200μL SCFA mix (SCFA group), once a day alone as respective treatment control. Dose and route of administration of SCFA mix was based on previous studies^20, 29, 30^. At the termination of the study, TRPA1-associated nociceptive behavioral assays (tail flick test and eye-wipe test) for loss of nociceptive responses was performed to ensure desensitization (Day-14). To study the effect of CPZ administration on neuronal TRPV1 activity, eye-wipe test (0.02% w/v capsaicin solution (Cat. No. 360376; Sigma Aldrich) and tail flick test (in water kept at 42°C) were performed at termination of the study. Replicates for each animal were taken using both eyes separately. Plan of experiment is explained in Fig. 1.

### 4.3. *In-vitro* gut permeability assay using FITC loaded solid-lipid nanoparticle

#### Formulation of FITC loaded solid-lipid nanoparticle

The fluorescent nanoparticle was prepared according to Ahmad *et al*.^31^. Briefly, 140 mg of glyceryl monostearate (GMS) (Cat. No. G0085; TCI Chemicals) and 5 mg of fluorescein isothiocyanate (FITC) (Cat. No. F0784; TCI Chemicals) were melted above 10°C of their melting points. The aqueous surfactant PF-127 (Cat. No. P2443; Sigma Aldrich) 2% w/v (200 mg) was dissolved in 10 ml of distilled water at same temperature. The aqueous phase was incorporated drop-by-drop into oil phase over constant stirring of 1500 rpm and kept for 1 hour, followed by sonication using probe sonicator for 10 minutes (5 seconds each on and off cycle at 40 Hz frequency). FITC-nanoparticles were lyophilized and dispersed in milli-Q water prior to administration.

#### *In-vitro* gut permeability assay

Determination of colon permeability was assessed as previously reported method^32^ with some modification. In brief, distal colon (2 cm) was excised and carefully flushed with normal saline, after that one end of the colon was ligated with the silk thread and 70μL of FITC loaded nanoparticles was filled in to the colon and the other end of the colon was also secured with silk thread. This assembly was then placed in a beaker filled with 150 ml of PBS buffer (pH – 7.4) and stirred at mild constant RPM (200 rpm). 1 ml of sample was withdrawn at pre-defined time intervals (0, 15, 30, 45, 60, 75 and 90 min) and replaced with fresh 1 ml of PBS to maintain sink condition. The withdrawn samples were analysed and imaged for fluorescent intensity by using IVIS Lumina Series-III (Perkin Elmer, USA) and analyzed using living image 5.0 software (Perkin Elmer, USA)

### 4.4. Gene expression analysis in DRGs & NanoString nCounter multiplex gene expression assay

Total RNA was extracted from dorsal root ganglion (DRGs) using TRIzol RNA extraction reagent-based method. Isolated RNA was transcribed into cDNA using Revert Aid cDNA synthesis kit (Cat. No. K1622; Thermo Fischer Scientific). The synthesized cDNA was then used to perform gene expression of specific markers of peptidergic sensory neurons^13^ by qRT-PCR using SYBR Green (Cat. No. L001752; Bio-Rad Laboratories). The conditions were at 95 °C for 2 min, followed by 40 cycles of annealing and elongation at 60 °C for 30 s and denaturation at 95 °C for 5 s. Analysis of relative change in gene expression was done using 2^-ΔΔCt^ method^33^. Ct values were normalized to β-actin (housekeeping) gene, and values were expressed in the terms of fold change with reference to control.

For transcriptional studies in colon, a customized Nanostring nCounter GX CodeSet gene expression panel (ILS_Bishnoi/C8788X1; NanoString Technologies, USA), comprising genes of goblet cell differentiation (*Cdx2*, *Dll1, Dll4, Foxa1, Foxa2, Foxa3, Klf1, Prkd1, Tff3, Vamp8*) / tight junction protein (TJPs) (*Cdh1, Cldn1, Cldn4, Occ, Zo1, Zo2*) / mucin production (*Muc15, Muc2, Muc20, Muc3, Muc4, Muc5b*) / glycosylation enzyme (*B4galnt1, B4galnt2, C1galt1, C3gnt1, Fut2, Galnt1, Galnt2, Galnt3, St3gal4*) / immune response (*Defb2, Dmbt1, Fcgbp, Nlrp3, Nod2, Reg3g, Src3, Tnfα, Xbp1*)/ 4 housekeeping genes (*Actb, B2m, Gapdh, Ubc*) was used. 50ng of extracted RNA sample was hybridized with unique probes using NanoString nCounter prep station and placed into the cartridge (NanoString Technologies, USA). NanoString nCounter digital analyzer (NanoString Technologies, USA) was employed for detection and counting of hybridized probes. The data was analyzed using nSolver 4.0 software (NanoString Technologies, USA).

### 4.5. Histology and immunohistochemistry Analysis

Ileum, proximal and distal colon were fixed in Carnoy’s solution (60% ethanol+30% chloroform+10% glacial acetic acid) and stored in the same until processed. Fixed tissues were serially dehydrated, cleared and embedded using gradient ethanol, xylene and molten paraffin respectively. 5-μm-thick sections were prepared, deparaffinized in xylene and rehydrated in gradient ethanol. For staining, Hematoxylin-Eosin (H&E) (Cat. No. S034, S007; Hi-media Laboratories) and Alcian blue (AB) (Cat. No. B8438; Hi-media Laboratories) were used for 30s, 15s and 15 min respectively. Slides were mounted using DPX mounting medium (Cat. No. GRM655; Hi-media Laboratories, India). Images were captured at 10X on bright field microscope (Leica CTR6, Leica Biosystems, Germany) and thickness of mucus layer was measured using Imagescope (Version 12.1.0.5029, Aperio Technologies Inc.) manually. For high iron diamine (HID) staining for sulfomucins, rehydrated tissue sections were stained with HID solution for 18 h at room temperature in the dark followed by washing with distilled water for 3 times. The sections were then stained with alcian blue for 30 minutes, washed 3 times with distilled water and mounted with DPX mounting medium. Images were captured at 20X on bright field microscope (Leica CTR6, Leica Biosystems, Germany).

For immunohistochemistry, 5-μm rehydrated sections were treated for antigen retrieval using citrate buffer (pH-6.0), blocked in 2.5% goat serum for 1 h followed by incubation in primary antibodies, Zona occludens-1 (*Zo1*)- (1:200 prepared in 2.5% goat serum, Cat. No. ab216880 Abcam), Occludin (*Occ*) (1:200 prepared in 2.5% goat serum, Cat. No. ab216327 Abcam), Claudin-1 (*Cldn-1*) (1:200 prepared in 2.5% goat serum, Cat. No. ab15098; Abcam) for overnight at 4°C. Finally, the sections were incubated with Alexa fluor 488 (1:1000 prepared in goat serum; Cell Signaling Technology, USA) secondary antibody for 1 h at room temperature. DAPI stain was used to stain nuclei. Images were taken under confocal microscope (Carl Zeiss LSM-880, Carl Zeiss-Germany) and all the images were captured at 40x magnification. Images were analyzed using Image J software (Public domain software, NIH, USA).

### 4.6. Scanning Electron Microscopy

For Scanning Electron Microscopic (SEM) analysis, small fragments of distal colon were fixed in 2.5% glutaraldehyde solution (Cat. No. G5882; Sigma Aldrich) in PBS (pH 7.4) for 6 h at 4°C. The tissues were washed with PBS, post-fixed in 1% osmium tetroxide (Cat. No. 201030; Sigma Aldrich) in PBS (pH 7.4) for 2 h at 4°C. Tissues were washed with PBS and dehydrated through the serial application of increasing concentration of acetone (Cat. No. 20003; SDFCL) viz. 30%, 50%, 70%, 90%, 95%, 100% to remove water at 4°C for 30 min period. Samples were air dried and spur-coated with gold (10 nm). Images were acquired using Apreo S SEM (Thermofischer Scientific, USA).

### 4.7. Analysis of microbial abundance in cecal content

DNA was isolated from 100 mg cecal content, using Nucleospin DNA Stool Genomic DNA Purification Kit (Cat. No. 740472; Macherey-Nagel, Germany), as per manufacturer’s instruction. DNA was quantified on spectrophotometer (Nanodrop, Thermo Scientific, USA) and run on 0.8% agarose gel electrophoresis for integrity. qRT-PCR was performed as per protocol detailed in section 4.4 using bacterial primers (Table 1). Total bacteria (TB) was used as internal control.

**Table-I:**
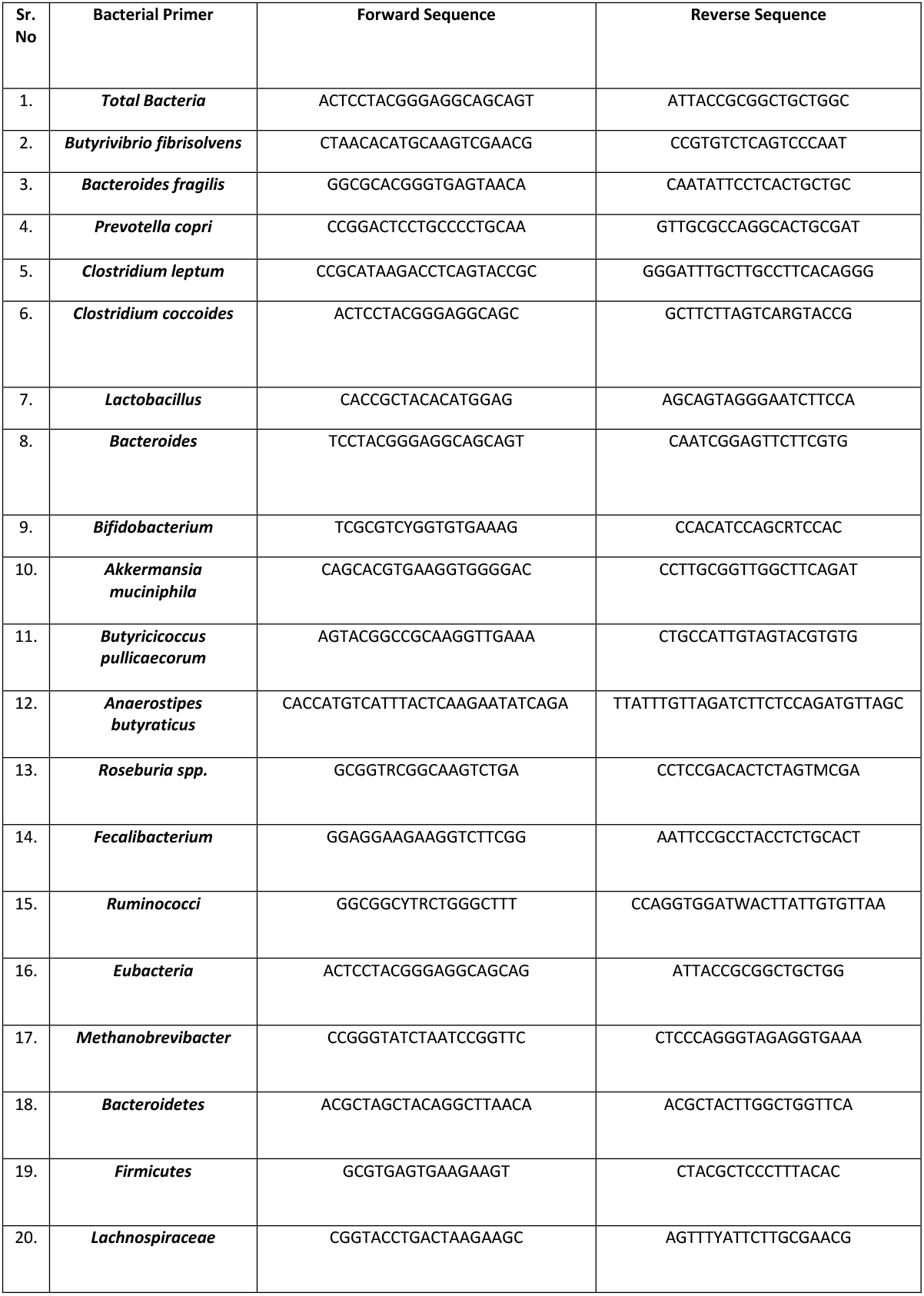
List of bacterial primers.

### 4.8. Metabolome profiling of cecal content

Gas chromatography-Mass spectrophotometry (GC-MS) was employed for metabolite analysis. Briefly, 80mg of cecal content was treated with methanol, chloroform and water (in 2.5:1:1 ratio) at 4°C, overnight with shaking. Samples were centrifuged (3000 rpm) and supernatant was lyophilized. For derivatization, lyophilized samples were treated with 90μL N, O-bis(trimethylsilyl)tri-fluoroacetamide (BSTFA), 1% trimethylchlorosilane (TMCS) (Cat. No. B-023; Merck & Co) and 30μL of pyridine (Cat. No. 029707; Central Drug House, India) and incubated for 4 h at 70°C. The samples were diluted up to 500μL with dichloromethane (DCM) (Cat. No. 508705; Central Drug House, India) and 5μL was injected in to GC-MS through automatic sampler. Agilent 7870A chromatograph equipped with an Agilent triplequadrupole 7000 mass spectrometer was employed for performing analysis. A DB-5 column (Agilent, 60 m x 250 μm ID x 0.25μm) was used for separation derivatized molecules before reached to MS chamber for their efficient identification. Column oven temperature was initially set at 80°C for 2 min, then increased to 140°C at 10°C/min, and then increased from 140 to 250°C at 5 °C/min. The helium carrier gas flow was set at 1.5 mL/min in spitless mode. For MS, the source temperature was set at 230°C, electron ionization was set at −70eV, the scan range was kept at 30-500 *m/z*, the gain factor was set 10 and thermal auxiliary temperatures 1 and 2 were set at 150 and 280°C respectively. Mass fragmentation of each peak of MS spectrum was matched with the NIST-MS library (version 2.0) for identifying metabolites

### 4.9. *In vitro* mouse fecal batch fermentation with CPZ and relative bacterial abundances

Slurry from freshly collected fecal samples (10%w/v) was prepared in 1X PBS containing cysteine with homogenisation. Fecal batch fermentation was done with 531μM CPZ and glucose (1% w/v) (Cat. No. MB-037; Himedia Laboratories, India) for 48 h. Basal growth media of the following composition was used (composition (g/l) peptone water 2g/l, yeast extract 2g/l, NaCl 0.1g/l, K_2_HPO_4_ 0.04g/l, KH_2_PO_4_ 0.04g/l, MgSO_4_.7H_2_O 0.01g/l, CaCl_2_.6H_2_O 0.01g/l, NaHCO_3_ 2g/l, Tween 80 2ml, hemin 0.02g/l, vitamin k1 10μl, cysteine HCl 0.5g/l, pH 7.0) (Himedia Laboratories, India). Initial pH was maintained at 7. Bacterial DNA extraction was done after 48h of fermentation using Nucleospin DNA Stool Genomic DNA Purification Kit (Macherey-Nagel, Germany) and relative abundance of selected gut bacteria was determined by qRT-PCR.

### 4.10. Statistics

Data was analysed using Graphpad Prism 8 software (Graphpad, USA). All the data is presented as mean ± SEM. Student unpaired t-test was employed to assess the statistical differences between the groups. P<0.05 was considered significant in all data. The principal component analysis was carried out using the adegenet (version 1.3-1) package in R^34^, while the correlation analysis (using pearson correlation coefficient) was carried out using the corrplot package in R.

## Acknowledgement

Authors would like to thank Department of Biotechnology (DBT), Government of India for research grant given to National Agri-Food Biotechnology Institute (NABI), Dr. Mahendra Bishnoi & Dr. Kanthi Kiran Kondepudi. We acknowledge Department of Biotechnology (Ramalingaswami Re-entry fellowship) & Department of Science, and Technology (DST-INSPIRE Faculty program) for providing grant to Dr. Ravindra Pal Singh and Dr. Sivasubramanian Rajarammohan respectively. We acknowledge DBT and University Grant Commission (UGC) for providing fellowship to Mr. Vibhu Kumar, Mr. Vijay Kumar and Ms. Kirti Devi. Authors are grateful to Dr. Arnab Mukhopadhyay and National Institute of Immunology (NII) for providing facilities and his team for helping to carry out nCounter experiment and analysis.

## Authors Contribution

VK, MB, KC and KKK generated hypothesis, conceptualized and designed the experiments. VK, KD and AK performed the experiments. VK, ViK, SR and MB conducted the analysis of the experiments. VK, ViK and MB wrote and edited the manuscript. RK, RPS, KKK and KC edited & reviewed the manuscript. KKK and MB lead contact, funding acquisition.

**Supplementary figure-1: Effect of CPZ rectal administration on TRPA1-induced nociceptive behavior and peptidergic sensory neuron markers on DRGs.**

A) AITC-induced eye wipe test and test for tail withdrawal latency at 4°C on day-7. B) AITC induced eye wipe test and test for tail withdrawal latency at 4°C on day-14. C) Capsaicin-induced eye wipe test and tail withdrawal latency test at 42°C on day-14. D) Representative image for classification of TRPA1-expressing colonic sensory neurons. E) Gene expression for DRG-associated makers for peptidergic sensory neurons. Mice were divided into two groups - Control (administered vehicle rectally) and CPZ (administered 531μM CPZ rectally). Treatment was given for 2 weeks. Desensitization of TRPA1 nociceptive neurons was confirmed with AITC (1mM) - induced eye wipes and Tail withdrawal latency test at 4°C on day-7 and day-14 (n=6). Desensitization of TRPV1 sensory neurons was confirmed with Capsaicin (0.02% w/v) - induced eye wipes and Tail withdrawal latency test at 42°C on day-14. After sacrifice, DRGs were harvested, RNA was isolated form DRGs and gene expression was performed for the markers of colonic peptidergic sensory neurons. All data is represented as mean ± SEM. Intergroup variations were assessed using Student unpaired t-test. *p<0.05, **p<0.01, *** p<0.001 versus Control.

**Supplementary figure-2: Effect of CPZ rectal administration on mucin glycans and genes involved in mucus glycosylation**

A) Metabolite profiling of cecum content for mucin glycans using GC-MS. B) Expressional changes in genes involved in mucus glycosylation. Mice were divided into two groups - Control (administered vehicle rectally) and CPZ (administered 531μM CPZ rectally).

Treatment was given for 2 weeks. Post-sacrifice, the cecum content was collected, weighed and treated overnight with methanol, chloroform and water in 2.5:1:1 ratio at 4°C (n=3). Methanol and chloroform were evaporated and sample was lyophilized. The samples were lyophilized and derivatized using BSTFA+1%TMCS, pyridine (70°C, 4 h). Following, the sample was diluted with DCM and run in GCMS. For gene expression studies, RNA was extracted from colon tissues and gene expression was performed for mucus glycosylation genes using Nanostring nCounter multiplex gene expression assay (n=6). All data is represented as mean ± SEM. Intergroup variations were assessed using Student unpaired t-test. *p<0.05, **p<0.01, *** p<0.001 versus Control.

**Supplementary figure-3: Direct effect of CPZ on *in vitro* fermentation of gut microbiota** A) Relative abundance of Gram -ve bacteria (after 48 h of fecal fermentation with or without 531μM CPZ treatment). B) Relative abundance of Gram +ve bacteria during *in vitro* fecal fermentation bacteria (after 48 h of fecal fermentation with without 531μM CPZ treatment). Briefly, fresh feces obtained from 12 normal C57BL/6J mice were pooled and a slurry of 0.1mg/ml was prepared in PBS. Cultures with 10X slurry dilution were prepared for: Control (treated with 1% w/v glucose) and treatment (treated with 531μM CPZ) groups (n=3). The cultures were incubated at 37°C under anaerobic conditions. After 48 h, the bacterial DNA was isolated from the culture using commercial kit. The DNA was subjected to qRT-PCR with primers of various bacterial genera. All data is represented as mean ± SEM. Intergroup variations were assessed using Student unpaired t-test. *p<0.05, **p<0.01, *** p<0.001 versus Control.

**Supplementary figure-4: Interaction between microbial population, metabolites and genes expression of Control and CPZ groups.**

A) Principal component analysis (PCA) plot for gut microbiota interactions between Control and CPZ treated group. B) PCA plot for interaction between metabolites in Control and CPZ group. C) PCA plot for interaction between gene expression changes in Control and CPZ group. D) Correlation matrix between parameters assessed in the Control and CPZ group. Both PC and correlation analysis were carried out using the adegenet and corrplot packages in R.

**Supplementary figure-5: Interaction between microbial population, metabolites and genes expression of SCFA and CPZ-SCFA groups.**

A) Principal component analysis (PCA) plot for gut microbiota interactions between SCFA (Control-SCFA) and CPZ-SCFA treated group. B) PCA plot for interaction between metabolites in Control-SCFA and CPZ-SCFA group. C) PCA plot for interaction between gene expression changes in Control-SCFA and CPZ-SCFA group. D) Correlation matrix between parameters assessed in the Control-SCFA and CPZ-SCFA group. Both PC and correlation analysis were carried out using the adegenet and corrplot packages in R.

