## Supplementary figures and images for "Colon Innervating TRPA1 Positive Nociceptors Influence Mucosal Health In Mice"

### Supplementary Figure-1

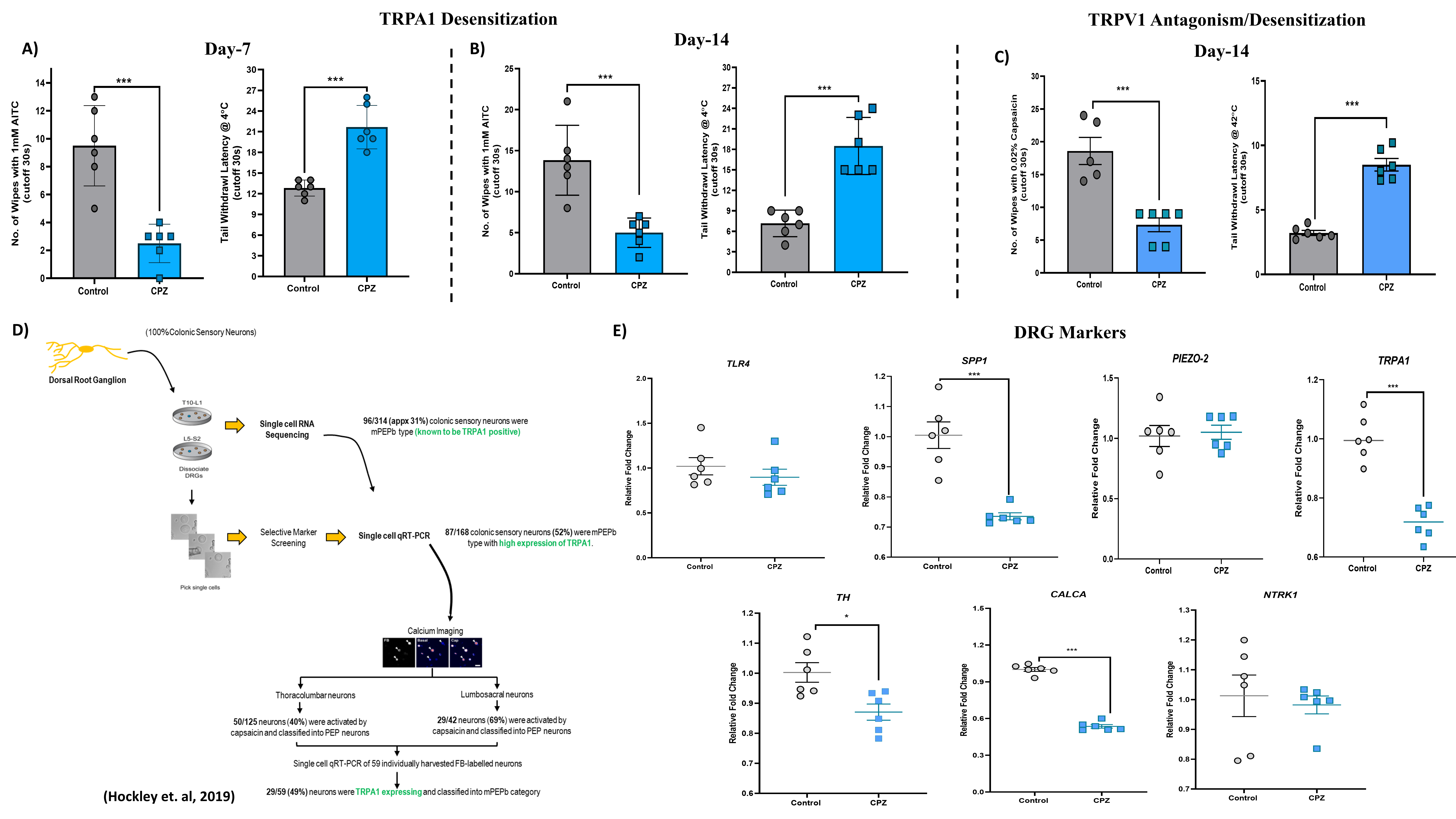

### Supplementary Figure-2

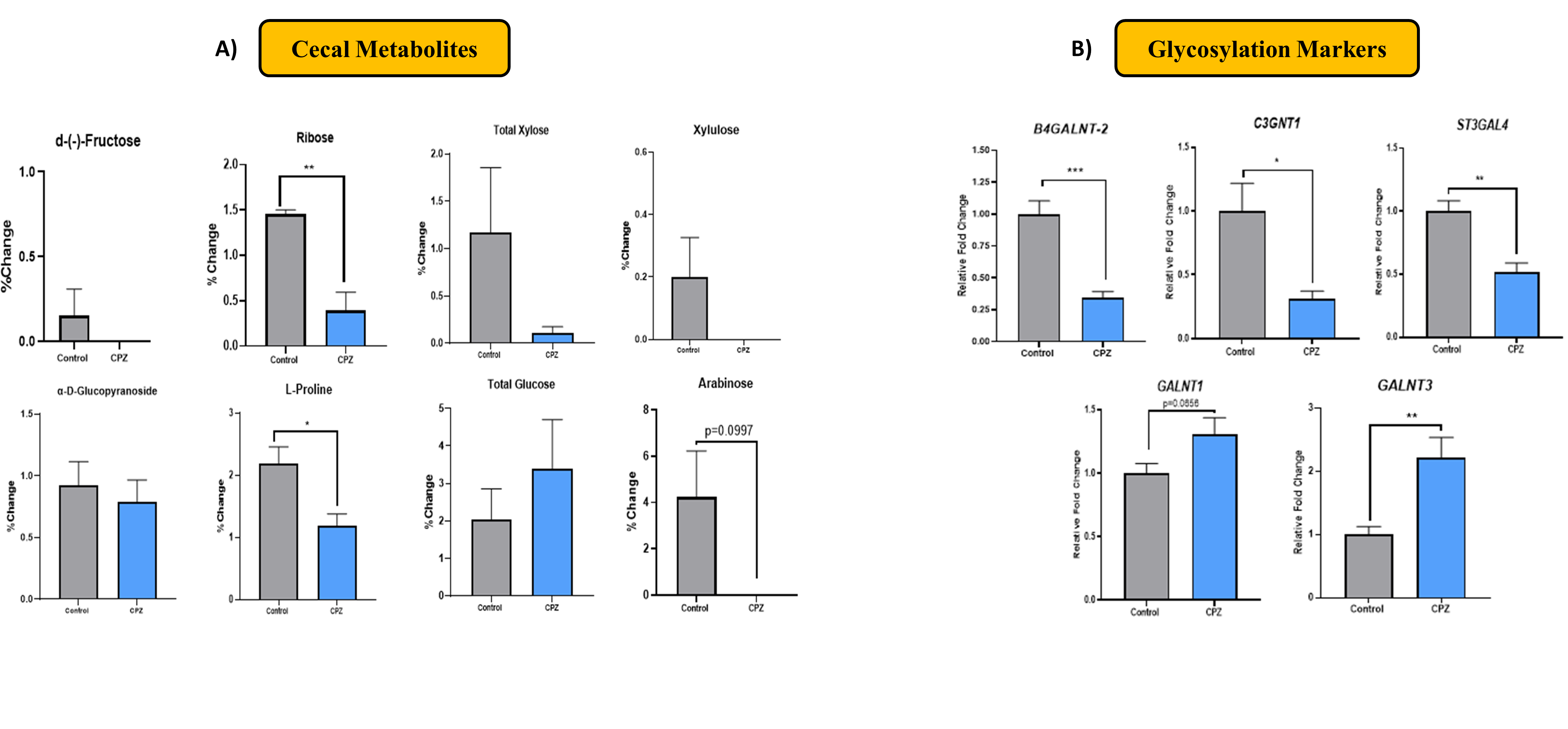

### Supplementary Figure-3

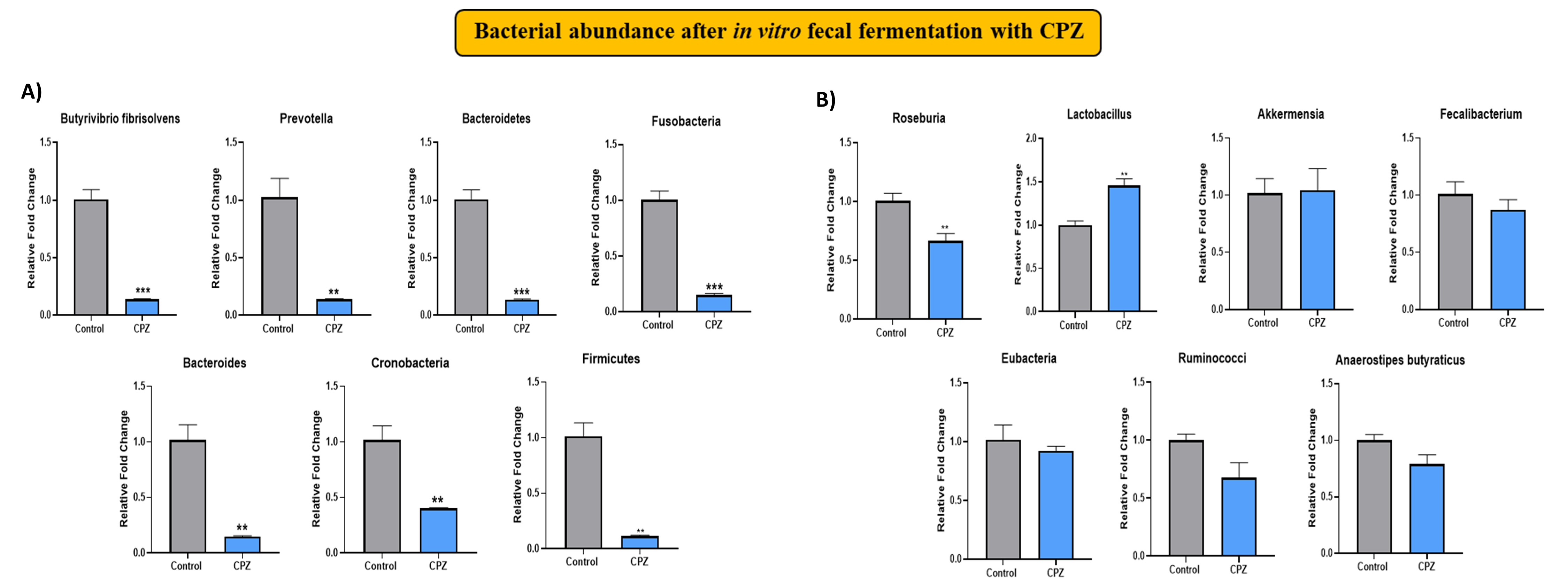

### Supplementary Figure-4

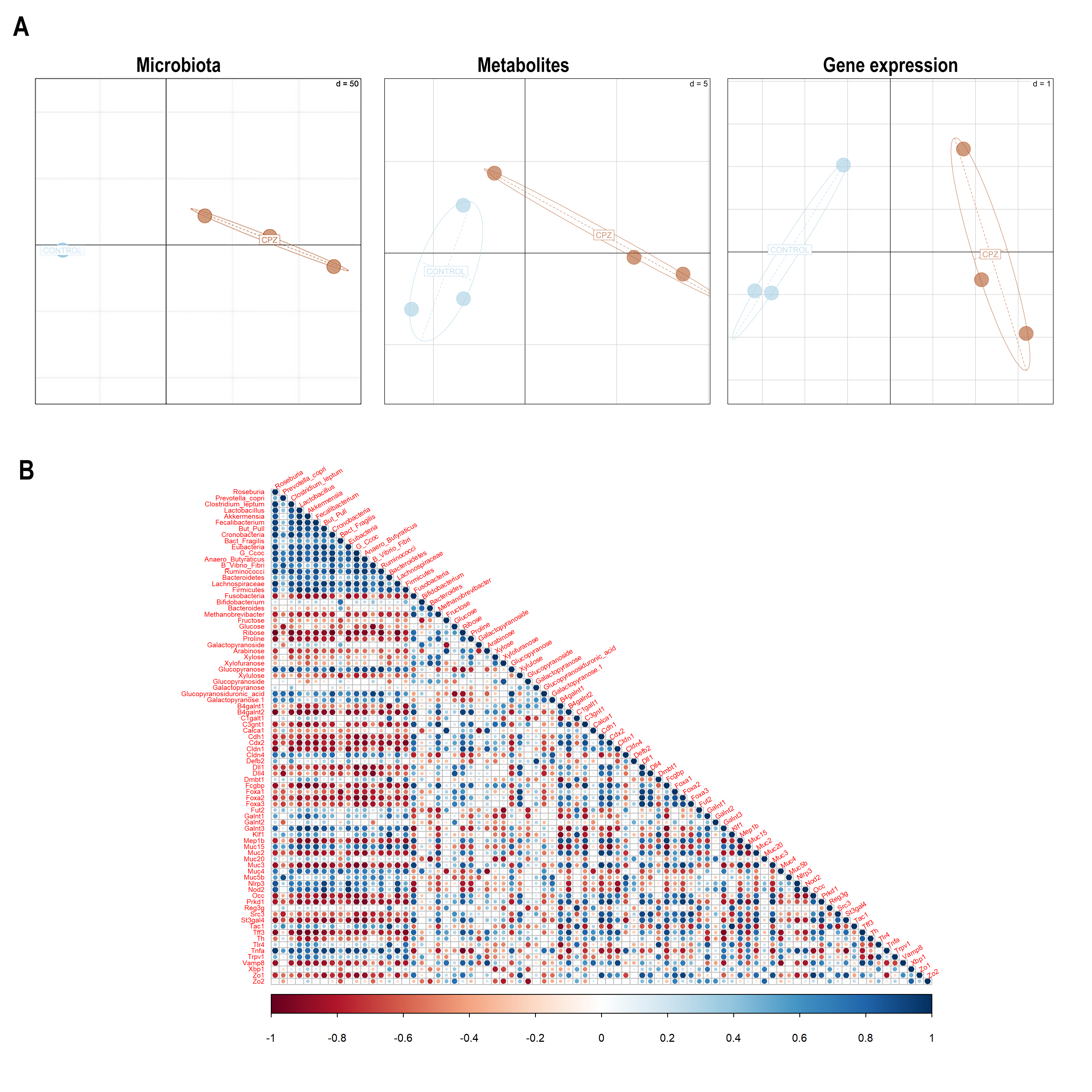

### Supplementary Figure-5

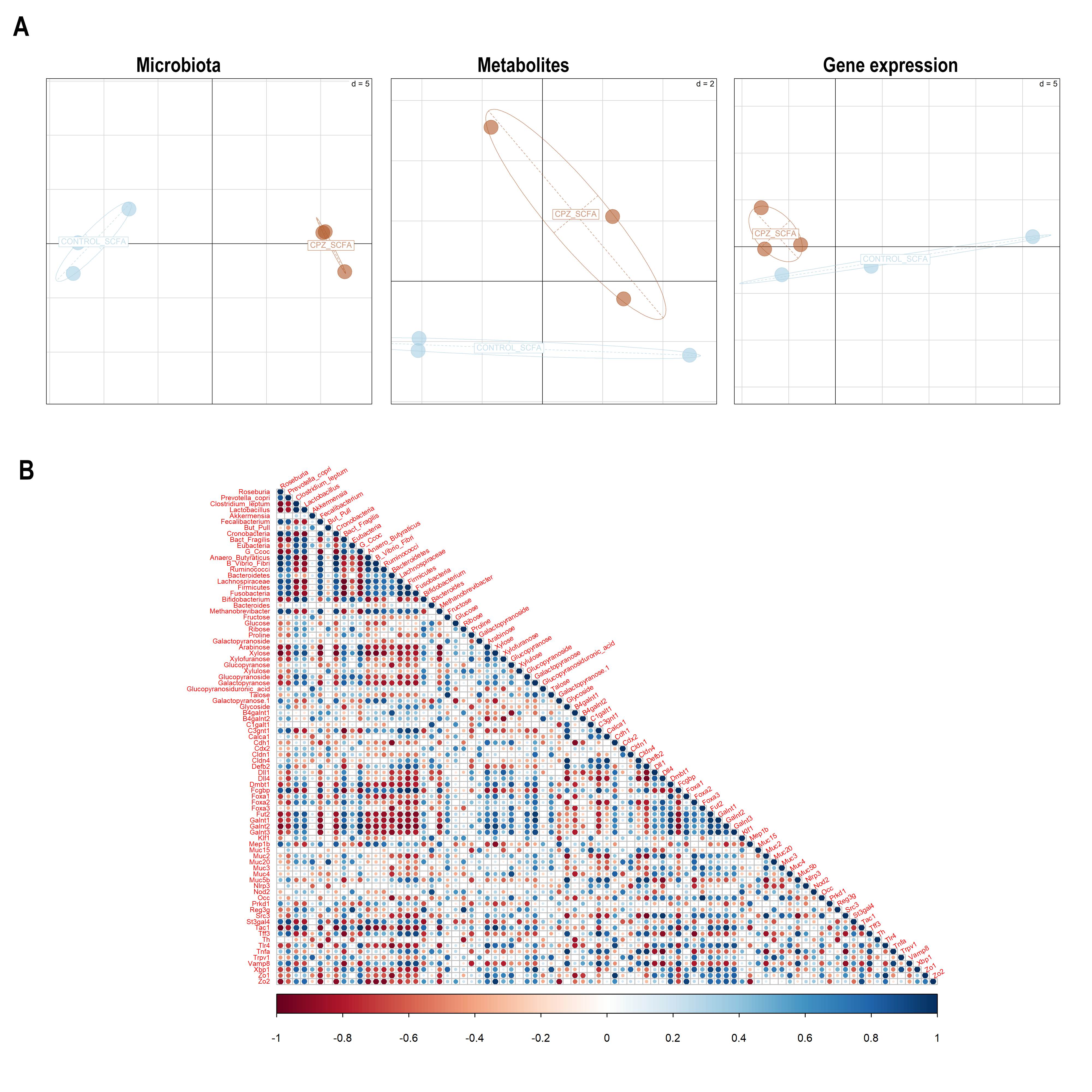
